# Three-dimensional Super-resolved Imaging of Paraffin-embedded Kidney Samples

**DOI:** 10.1101/2021.09.02.458679

**Authors:** David Unnersjö-Jess, Amer Ramdedovic, Martin Höhne, Linus Butt, Felix C. Koehler, Roman-Ulrich Müller, Peter F. Hoyer, Hans Blom, Bernhard Schermer, Thomas Benzing

## Abstract

Diseases of the glomeruli, the renal filtration units, are a leading cause of progressive kidney disease. Assessment of the ultrastructure of podocytes at the glomerular filtration barrier is essential for diagnosing diverse disease entities, providing insight into the disease pathogenesis as well as monitoring treatment responses. New technologies, including superresolved nanoscopy and expansion microscopy, as well as new sample preparation techniques, are starting to revolutionize imaging of biopsy specimens. However, our previous approaches for simple and fast three-dimensional imaging of optically cleared samples are to date not compatible with formalin fixed paraffin-embedded (FFPE) tissue, impeding application in clinical routine. Here we provide protocols that circumvent these limitations and allow for three dimensional STED and confocal imaging of FFPE kidney tissue with similar staining and image quality as compared to our previous approaches. This would increase the feasibility to implement these protocols in clinical routines, as FFPE is the gold standard method for storage of patient samples.

## INTRODUCTION

The three-layered kidney filtration barrier is responsible for filtering 180 liters of primary urine daily. It consists of a fenestrated endothelium, a uniquely composed glomerular basement membrane and an epithelial layer of podocytes, forming interdigitating structures called foot processes (FP)^1^. Especially the visualization of FP is of high importance to assess glomerular health, since effacement (i.e. morphological broadening and shortening) of FP is seen in almost all types of glomerular disease. Due to the tiny dimensions of these structures (width 200-1000 nm, spacing ~ 50 nm) the standard imaging modality used to visualize FP has previously been scanning or transmission electron microscopy^2^. While powerful in resolution, electron microscopy has drawbacks and limitations when it comes to non-destructive and straight-forward three-dimensional imaging. Further, since electron microscopy produces grayscale contrast images of all constituents of the tissue, segmentation of structures of interest for further analysis can be challenging^3^. In optical microscopy however, tagging specific molecules with fluorescent dyes allows for visualizing only structures of interest while no signal is collected from other regions, thus substantially simplifying automated image segmentation. In the past decade, new optical imaging techniques have been developed for visualizing FP and other parts of the filtration barrier, such as the glomerular basement membrane (GBM)^4^. In 2016, it was, for the first time, shown that podocyte FP and the slit diaphragm (SD) can be resolved *in situ* in three dimensions using a combination of stimulated emission depletion (STED) microscopy and tissue clearing^5^. It was also shown in this study that FP pathology (effacement) can be quantified in terms of the SD coverage. Further development, by our group and others, has made it possible to automatically extract the SD coverage^6,7^ and a plethora of other quantitative parameters describing effacement^7,8^. Our group has recently shown that SD and FP morphometry can be extracted from fluorescence images in a completely automated manner by applying deeplearning segmentation, further establishing the quantitative advantages of optical microscopy for imaging FP and quantifying effacement^8^. Prior to this, we moreover published a straightforward 3D imaging and quantitative pathological analysis protocol for both mouse and human samples which can be finished for full biopsy diagnostics in only five hours^9^. All of the protocols previously developed by us are based on optically clear, thicker Vibratome sections of fresh, fresh-frozen or PFA fixed samples/biopsies. Other than the obvious advantage of three-dimensional capacity, which gives access to a much larger volume for imaging and analysis, these protocols have proven highly superior to non-cleared samples when it comes to achieving high labelling quality and contrast^5,9^. The focus of this study is to investigate how our clearing and swelling protocols perform with paraffin embedded tissue/biopsies. This will be of importance in many research or pathology labs, as paraffin embedding is a old standard of storing tissue for later analysis. We show that, by slightly modifying our protocols, super-resolved STED and confocal, three-dimensional imaging of podocyte morphology can readily be performed in samples that have been formalin fixed and paraffin embedded. This broadens the use of our fast and simple clearing and swelling protocols for renal research, but especially for accelerated use in clinical pathology.

## RESULTS

### Optical Clearing and Mild Swelling of Kidney Tissue

The general workflow is schematically shown in Fig. 1a. We used our recently published fast clearing and swelling protocol^9^ as a starting point, while adding a deparaffinization step prior to Vibratome sectioning. Samples were treated according to a slightly modified simple clearing protocol and podocyte substructure could be imaged using either confocal or STED microscopy with subsequent segmentation and morphological analysis (Fig. 1b-d). One of the most important advantages of our previously published protocols is the access to the depth dimension and we therefore wanted to explore if the depth dimension would be accessible also in FFPE tissue. To this end, pieces of kidney tissue were cut out of the paraffin block using a razor blade and were then de-paraffinized using standard de-paraffinization protocols, utilizing Xylene for dissolving paraffin followed by re-hydration in decreasing concentrations of ethanol in water. After this, samples were cut into 200 μm thick sections using a Vibratome. Samples were then de-lipidated using SDS, and different clearing times and temperatures were tried (Fig. S1). As compared to non-paraffinized PFA or formalin fixed samples, slightly longer clearing times and higher temperatures were needed for sufficient clearing (Fig. S1a-b). Of note is that the nephrin staining, the basis for later quantitative analysis, showed sufficient signal after only 15 min of clearing, whereas podocin needed longer clearing time for sufficient antigenicity (Fig S1a). Thus, if only nephrin is to be stained for, the protocol can be shortened. Standard antigen retrieval by boiling in Tris-EDTA was tried, resulting in slightly lower staining quality and less optical transparency, and we thus decided to keep the SDS clearing step (Fig. S1c). Based on the results in Fig. S1, we established an SDS clearing step of 2 hours at 80 degrees as a starting point. However, as with all clearing protocols (and also with standard antigen retrieval on thin sections), temperature and time need to be optimized based on tissue type, fixation time and conditions, different epitopes/antibodies etc. Our final protocol resulted in a linear swelling of around 25%, slightly lower than earlier reported in non-FFPE samples^9^ (Fig. S1d).

**Figure 1.**
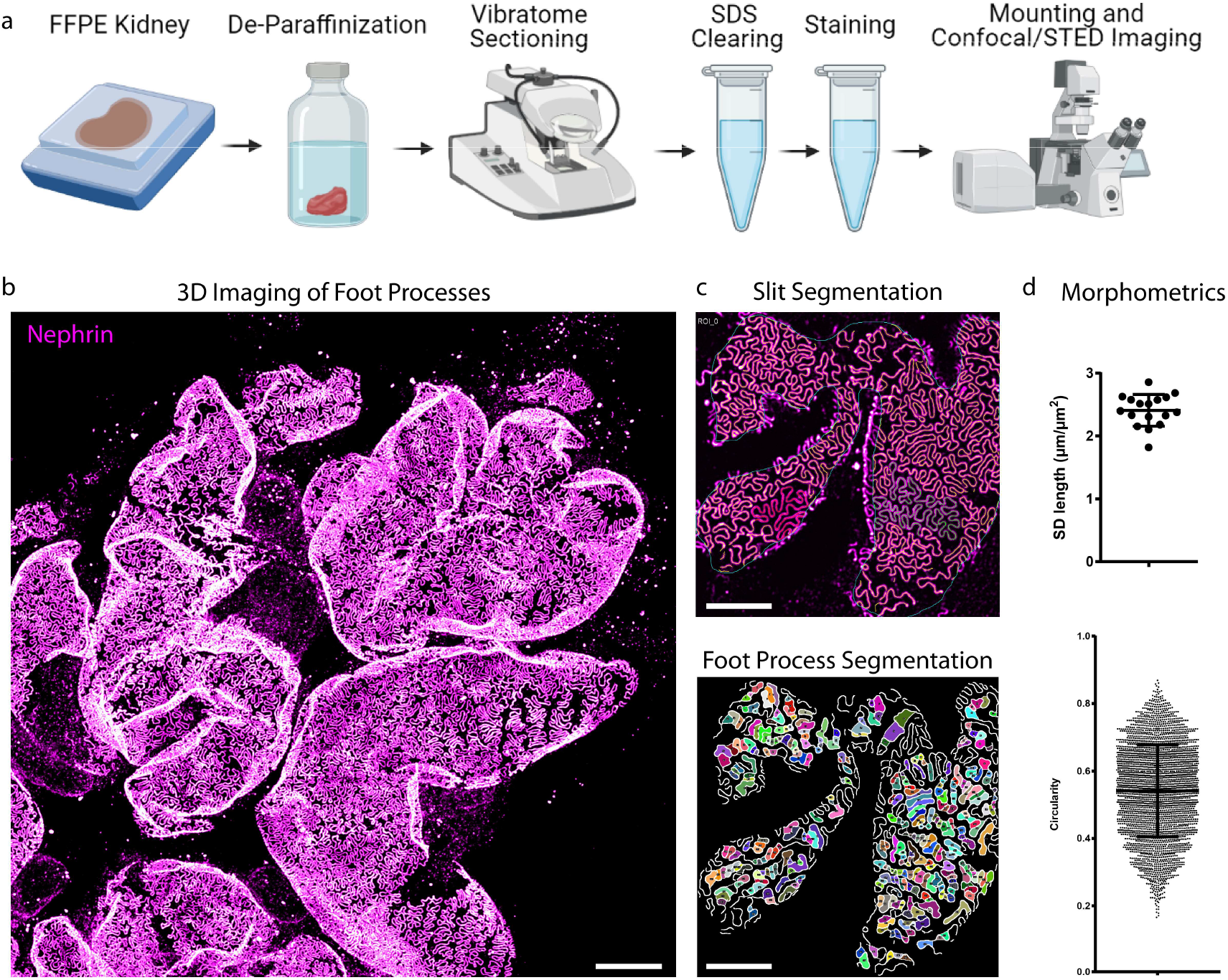
Fast sample preparation and imaging protocol for the visualization of foot processes in FFPE kidney tissue. (a) Overview of the sample preparation and imaging protocol. (b-d) This protocol allows for fast, high throughput confocal imaging of FP on glomerular capillaries (b), with semi-automatic segmentation of the SD and FP (c) and quantification of morphometry (d). The images and plots are illustrative examples, see figure 2–4 for detailed results. Scale bars 10 μm (b), 5 μm (c).

### Visualizing FP and the SD using STED and Confocal Microscopy

We next validated, by staining for podocin and nephrin, that FP in both mouse and human tissue could be clearly resolved using STED microscopy (Fig. 2a,d). Importantly, the expression of podocin on both sides of the slit could be resolved (Fig. 2b-c,e-f), which validates that the resolution achieved with this FFPE-optimized protocol is similar as compared to our previous, more tedious, STED-imaging protocol^5^. In our latest fast and simple protocol, we show that podocyte substructure can be resolved using confocal microscopy if samples are embedded in a fructose solution containing 4M urea, in order to induce a slight swelling of the sample^9^. We show that this approach can be applied also to FFPE tissue by demonstrating that FP can clearly be resolved in both a human and a mouse sample (Fig. 2g-h).

**Figure 2.**
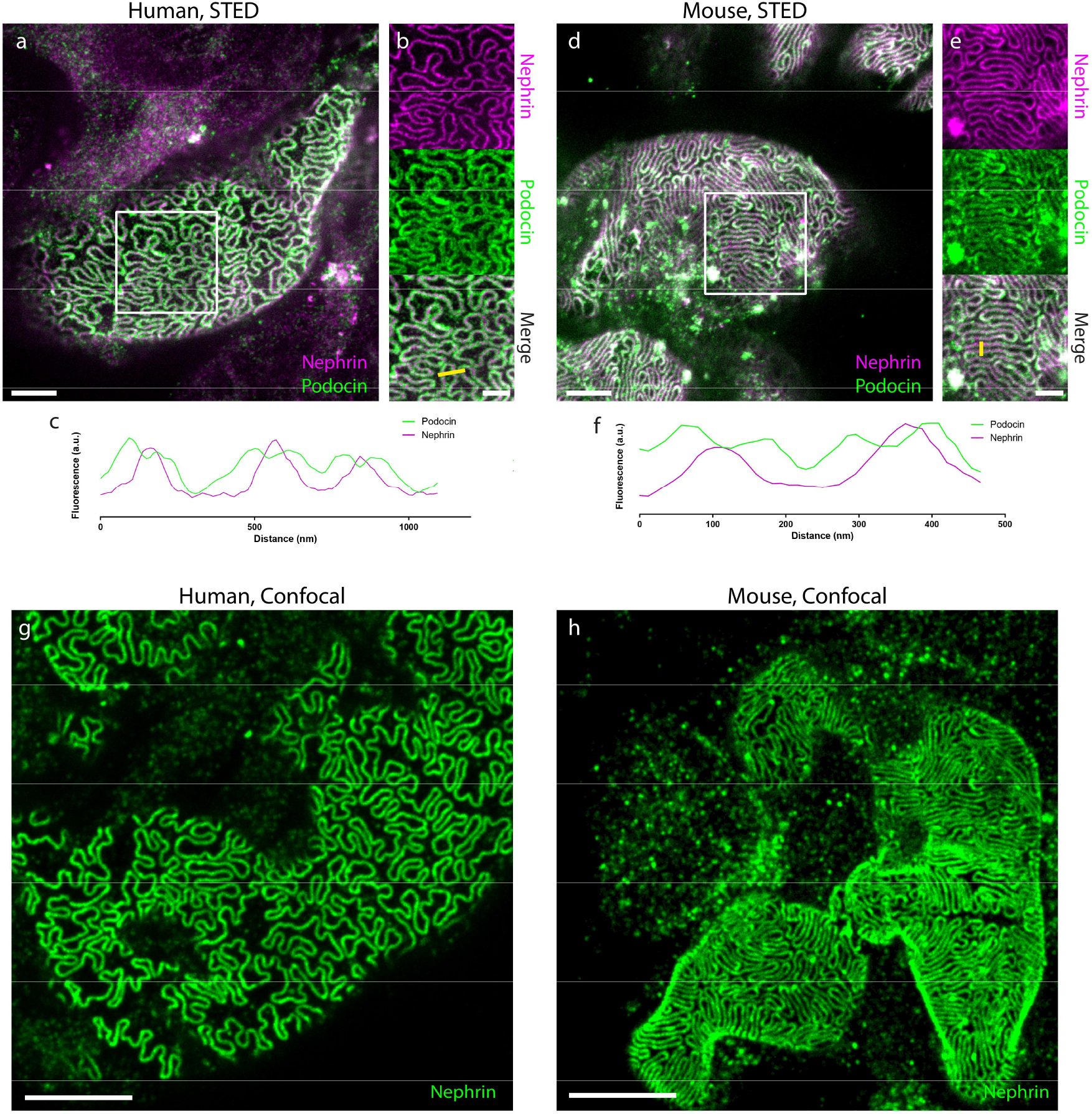
Validation of the protocol for imaging podocyte ultrastructure in human and mouse FFPE tissue samples. All samples were imaged using a Leica SP8 3X STED microscope with a 100X/1.4NA oil objective. All images are maximum intensity projections of ~ 2 μm thick z-stacks acquired at depths of 0-20 μm. Samples are stained for nephrin using either Abberior STAR-635P (a-e) or Alexa-488 (g-h) and for podocin using Atto-594 (a-e). Scale bars 2 μm (a, d), 1 μm (b, e), 5 μm (g, h). (a) A human control sample treated according to the protocol shows that FP are clearly outlined by staining for the SD. (b) Single channel views of the area outlined by a white rectangle in (a) shows that the two-sided expression of podocin on both sides of the SD can be resolved. (c) A line-profile along the yellow line in (b) shows that podocin is flanking the nephrin signal in three SDs with a separation of the podocin signal of around 80-100 nm. (d) Also in a wild-type mouse sample, FP are clearly outlined by staining for the SD. (e) Single channel views of the area outlined by a white rectangle in (d) shows that the two-sided expression of podocin on both sides of the SD can be resolved also in the mouse sample. (f) A line-profile along the yellow line in (e) shows that podocin is flanking the nephrin signal in two SDs, also here with a separation of the podocin signal of around 80100 nm. (g-h) Confocal images of human (g) and mouse (h) tissue, showing that foot process morphology can be resolved with diffraction-limited microscopy by using a 1.4 NA 100X objective with the green-emitting dye Alexa-488 and a confocal pinhole size of 0.3 airy units.

We next applied our validated clearing and swelling protocol to clinical samples. Occasionally, such as when retrieving samples from a biobank or a pathology lab, the only option is to get thin (<10 μm) paraffin sections on glass slides. Thus, we investigated if our protocol could be applied also to such samples. Applying a standard Tris-EDTA antigen retrieval step prior to staining with the simplified staining protocol used for cleared sections^9^ did not result in sufficient staining quality to resolve foot processes (Fig. S2a). Interestingly, we found that removal of the section from the glass slide using a razor blade and performing the staining free-floating, substantially increased staining quality (Fig. S2b). However, it was only after adding a 1-hour de-lipidation step at 80°C that sufficient staining quality could be achieved to resolve the SD by staining for podocin (Fig. S2c-d). Importantly, we did manage to achieve sufficient staining quality also in sections mounted on a glass slide by extending the Tris-EDTA boil to 40 minutes in length and by applying a more laborious standard immunohistochemistry staining protocol, using blocking with BSA/NDS and by adding several washing steps between and after antibody incubations (Fig. S2f). However, we were never able to resolve the SD when staining for podocin in sections still mounted on a glass slide (Fig. S2g). The results from Fig. S2 further establishes what has earlier been shown, that SDS lipidation of thick sections results in higher staining quality, and that this staining quality can be achieved also by using a drastically simplified staining protocol with no blocking and very little washing^5,9^.

### Quantifying Glomerular Nanoscale Pathology in FFPE Samples from Mice and Humans

We further went on to validate that, by applying our previously published analysis approaches, we could quantitatively describe pathological FP effacement in both humans and mice. To do this, we applied the protocol to FFPE tissue from a human patient with trans-associated mutations in TRPC6 and NPHS2, and to mice with trans-associated mutations in the NPHS2 gene (R231Q/A286V), both leading to an FSGS phenotype (Fig. 3). Using a previously published ImageJ macro^7^, we show that both the filtration slit pattern and individual foot processes could be segmented from images acquired using confocal microscopy (Fig. 3a,d). From these segmentations, the SD length per area (Fig. 3b,e) as well as FP circularity (Fig. 3c,f) could be extracted. The results for the mouse FSGS model go well in line with what has previously been published^7^, with a significantly lower SD length and a significantly increased FP circularity as compared to controls (Fig. 3e-f). A decrease in SD length was also observed for the human patient (Fig. 3b) whereas the increased circularity as seen in the mouse FSGS model could not be observed. Instead, a decrease in circularity was observed for the patient with genetic FSGS (Fig. 3c). Also for other morphometric parameters, except for FP area, results for FSGS mice and the FSGS human patient showed opposite significant differences (Fig. S2). One reason for this variability could be that the control human sample was taken from a very old patient (> 70 y/o) whereas the FSGS sample was taken from a young child (< 1 y/o), while mouse samples were age-matched. Moreover, we have recently shown that different disease types can present with different morphological effacement “fingerprints”, and thus, effacement patterns might differ between subtypes of FSGS^8^. Even though the segmentation and analysis works well in images acquired with confocal microscopy, we further demonstrate that, as a result of the higher spatial resolution, the ImageJ macros perform slightly better in images acquired with STED microscopy (Fig. S4). As part of the SD density macro, the user has to define the capillary surface area in which the analysis is to be carried out. This is not always straight-forward, especially in capillary regions with severe FP effacement, and might lead to erroneous results. We found that staining the GBM using a collagen-IV antibody enabled us to automatically and accurately define the region-of-interest (ROI) without user intervention (Fig. S5).

**Figure 3.**
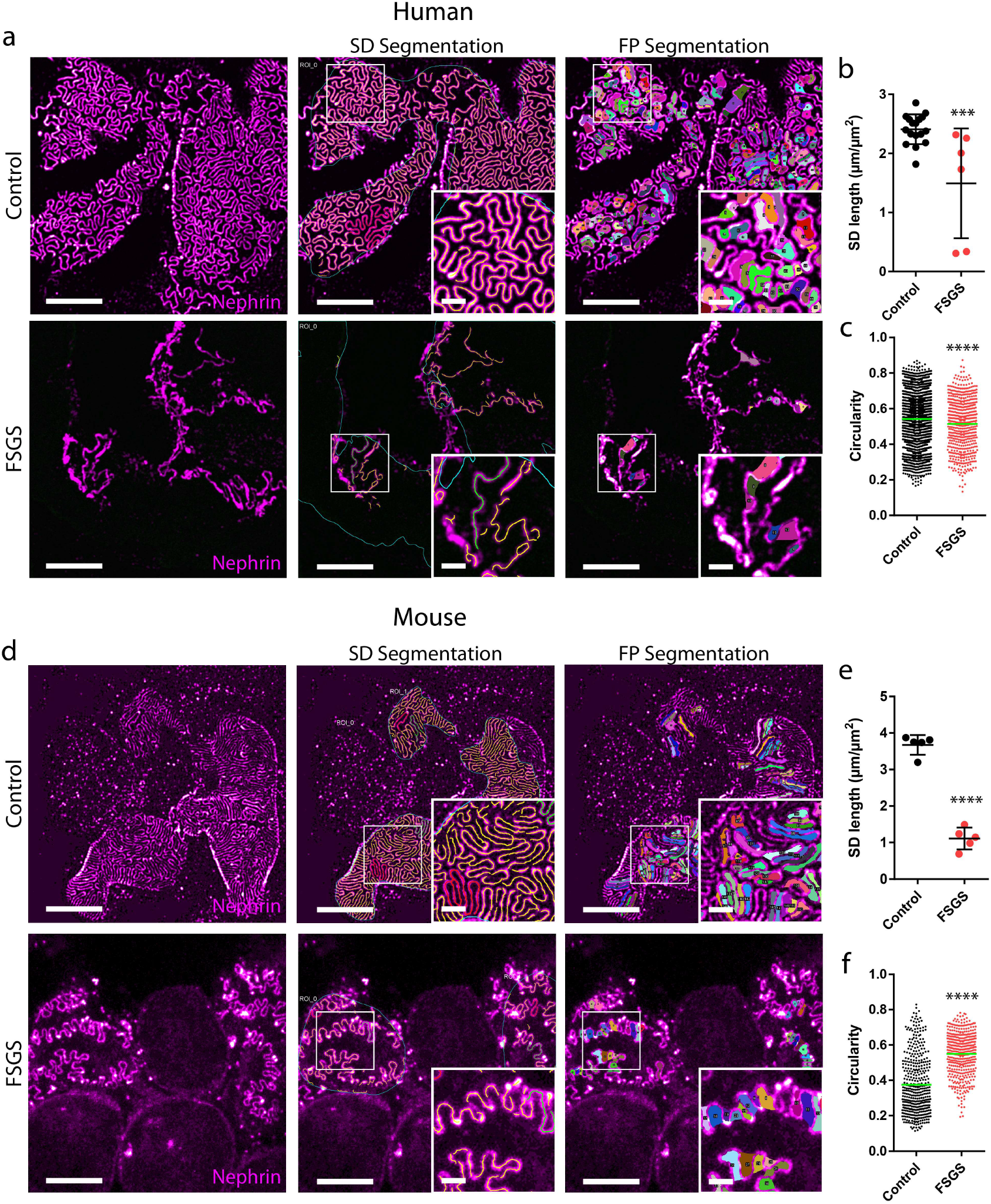
Imaging and segmentation/morphometry of the SD/FP pattern in healthy and diseased human and mouse FFPE tissue imaged with confocal microscopy. All samples were stained for nephrin using Alexa-488 and imaged using a 100X NA 1.4 oil objective with a confocal pinhole setting of 0.3 AU. Scale bars 5 μm and 1 μm (zoomed insets). (a) Maximum intensity projections of ~ 2 μm thick z-stacks showing the global slit pattern in a control human sample and a sample from a patient diagnosed with FSGS. Insets show zooms of the area marked with a white square showing effacement in higher detail. (b) SD length per capillary area in control and FSGS patients, showing a significant decrease for the FSGS patient. Each dot represents one image. Black line represents mean and error bars represent standard deviation. Two-tailed t-test, P = 0.001. (c) FP circularity score for Control and FSGS patients showing an unexpected but significant decrease in circularity score for the FSGS patient. Each dot represents one foot processes and the green line represents mean. Two-tailed t-test, P < 0.0001. (d) Maximum intensity projections of ~ 2 μm thick z-stacks showing the global SD pattern in a WT mouse sample and a sample from a mouse with genetic FSGS. Insets show zooms of the area marked with a white square showing effacement of FP in the FSGS mouse in higher detail. (e) SD length per capillary area in WT and FSGS mice, showing a significant decrease for the mutated mouse. Each dot represents one image. Black line represents mean and error bars represent standard deviation. Two-tailed t-test, P < 0.0001. (f) FP circularity score for WT and FSGS mice showing a significant increase in circularity score for the FSGS mouse. Each dot represents one FP and the green line represents mean. Two-tailed t-test, P < 0.0001.

### Three-dimensional Confocal and STED Imaging of Thick Samples

We have previously shown that our optical clearing protocols allow for both confocal and STED imaging deep inside intact kidney tissue^5,9,10^. Also with our FFPE-optimized protocol, we validate that thick samples can be imaged in three dimensions both at low and high magnification (Fig. 4a-c). By mixing the fructose embedding medium at a lower concentration, the refractive index can be freely adjusted to 1.45, perfectly matching a longer working distance glycerol objective lens. With this approach, confocal imaging is possible at depths of at least 120 μm (Fig. S6) in cleared FFPE samples, although the lower NA of the glycerol objective lens of 1.3 results in slightly lower lateral and axial resolution as compared to the 1.4 NA oil objective otherwise used. Although sufficient lateral resolution for resolving FP can be achieved with a confocal microscope, the z-resolution is still not sufficient to resolve FP when projecting x-z or y-z directions on capillaries (Fig 4d). However, by applying 3D STED, which increases the axial resolving power by a factor of 5, sufficient isotropic resolution is reached for resolving FP nanoscale morphology in any spatial orientation (Fig. 4e-g, Supplementary Movie 1). We have thus shown that our in-situ 3D imaging capacity shown with previous protocols is transferable to clinically stored FFPE tissue samples as well.

**Figure 4.**
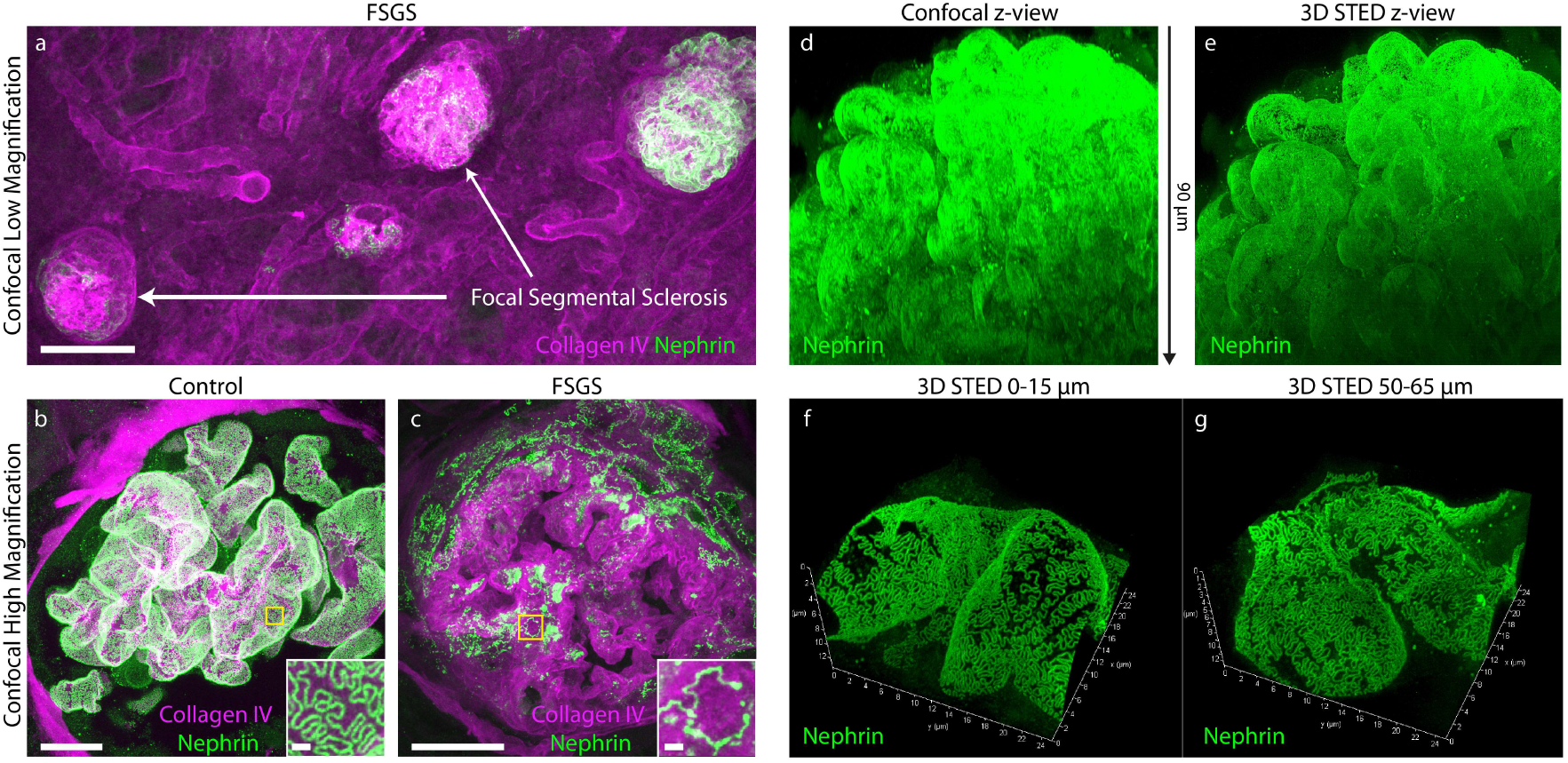
Three-dimensional confocal and STED imaging in human samples (same samples as in Fig. 3). All images acquired using a Leica SP8 confocal system using either a 20X NA 0.75 oil (a), a 100X NA 1.4 oil objective or a 93X NA 1.3 glycerol motCorr objective (d-c). Samples were stained for nephrin with either Alexa-488 (a-c) or Abberior STAR635P (a) Maximum intensity projection of ~ 200 μm thick z-stack in samples from the FSGS patient showing clear focal and segmental sclerosis of glomeruli (high collagen expression and low nephrin expression). Scale bar 100 μm. (b-c) Maximum intensity projections of ~ 100 μm thick confocal z-stacks of control (b) and FSGS (c) patients, showing apparent effacement of FP in a sclerosed glomerulus of the FSGS patient. Insets show zooms of areas marked with yellow squares. Scale bars 10 μm and 1 μm (zoomed insets). (d-e) Maximum intensity projections (x-z views) of confocal (d) and 3D STED (e) stacks of the same glomerulus. The z-depth is indicated with the arrow between the images. See Supplementary Video 1 for better visualization of the data. (f-g) Maximum intensity projections of 3D STED stacks of glomerular capillaries in the control patient, showing that FP can be clearly resolved at depths of at least 65 μm. Stacks are tilted to show the high z-resolution provided by the STED depletion in the z-dimension. Dimensions are indicated on the axes.

## DISCUSSION

We have in the last years shown that, by applying novel sample preparation technologies, superb staining quality can be achieved when imaging podocyte substructure in intact kidney tissue. Apart from higher signal-to-noise ratio and higher labelling density, the protocols also allow for three-dimensional visualization of FP morphology in health and disease. The protocols have proven to be superior for later semi-automatic and automatic segmentation to extract quantitative parameters describing podocyte morphology^7,8^. By modifying these protocols, we here show that also FFPE tissue can be treated according to our protocol to achieve similar imaging results as with samples not embedded in paraffin. We validated that also our previously developed image analysis/quantification strategies can be applied to FFPE samples, with correlating results. In FFPE tissue, we found that SDS de-lipidation acts not only to increase tissue transparency, but also as an antigen retrieval step, with increasing epitope availability upon increased clearing time. We further show that the increase in staining quality can be partially attributed to the fact that samples are stained in a free-floating manner, since an identical staining protocol applied to an FFPE section mounted on a glass slide resulted in substantially lower staining quality. However, as has also been demonstrated previously^6,11,12^, we demonstrate that it is possible to achieve decent staining quality also in sections mounted on glass slides by applying a longer antigen retrieval step and a more tedious labelling protocol with extensive washing and blocking. For maximal staining quality and effective resolution, FFPE tissue samples had to be cleared and stained in a free-floating manner. We thus show that, apart from the obvious disadvantage of having access to less in *situ* 3D volume for imaging, thin FFPE sections mounted on slides are sub-optimal for visualizing and segmenting podocyte substructure.

Taken together, findings in this study show, that our fast and simple clearing protocols can be applied also to FFPE tissue blocks, which has its main impact when it comes to the study of human clinical samples. In most clinical pathology labs, it is standard procedure to embed biopsies in paraffin before thin sectioning and histological evaluation. Thus, the threshold for implementing our clearing protocols in a clinical pathology workflow is drastically lowered.

It should be pointed out that the long de-paraffinization protocol applied here to tissue chunks with thickness of several millimeters can be drastically shortened when de-paraffinizing smaller samples, such as human biopsies. It should also be apparent that paraffinization and de-paraffinization of tissue is still laborious and time-consuming, and there is thus no benefit in applying it for the purpose of imaging FP using optical microscopy, if one has the choice. Moreover, our recently shown automatic multi-parametric morphological analysis of filtration structure morphology, based on deep learning, gives additional information regarding differential patterns of effacement across different diagnoses^8^. By applying this analysis strategy to images provided by the protocol in this study with just classical confocal imaging, a faster and completely automated and bias-free evaluation of effacement in biopsies could possibly be implemented in clinical pathology in the future. Finally, it should be mentioned that other protocols exist for imaging foot processes in FFPE samples, but all of these superresolution approaches are applied to thin sections, with their aforementioned disadvantages^6,13^. In addition, we have previously shown that all of these published protocols require more reagents and user interactions as compared to the simple protocol applied in this study which, again, impedes feasible transfer of new pathology approaches to the clinic.

## MATERIALS AND METHODS

### Mouse and Human Kidney Tissue

All mouse experiments were approved by the State Office of North Rhine-Westphalia, Department of Nature, Environment and Consumer Protection (LANUV NRW, Germany) and were performed in accordance with European, national and institutional guidelines. Mice of 100% C57BL/6N background were used. After anesthesia with Ketamine and Xylazine, mice were euthanized by cardiac perfusion with Hank’s Balanced Salt Solution (HBSS) and fixated as described below.

Patient material was obtained from one child suffering from Steroid-Resistant Nephrotic Syndrome due to compound-heterozygous point mutations in TRPC6 and NPHS2. Control human tissue was collected from a patient that was nephrectomized due to renal carcinoma. Tissue sample was dissected from the non-tumorous pole of the kidney and showed normal histological picture in routine histological examination. All procedures were approved by the Ethics Commission of Cologne (DRKS00024517) and conducted in accordance with the declaration of Helsinki. Patients or the patient’s parents gave informed consent.

### Mouse Model for FSGS

Mice with two compound-heterozygous point mutations, Pod^R231Q/A286V^, were generated as previously described^7^. Mice were sacrificed at 20 weeks of age as stated above.

### Fixation and Paraffin Embedding

Mouse kidneys were fixed in 4% PFA in 1X PBS at room temperature for 1-3 hours. Healthy kidney tissue of human control patients was obtained from the healthy contralateral pole of the renal carcinoma and was fixed in 4% PFA in 1X PBS overnight. Mouse and human tissue was then paraffin embedded using standard procedures.

### De-paraffinization

~1-2 mm thick pieces of cortex was cut from the paraffin block using a scalpel. Alternatively, samples were sectioned to 10 μm thickness using a microtome and mounted on glass slides. Paraffin was dissolved by incubating the samples in Xylene over night (3*5 min for thin sections). After this the samples were re-hydrated by incubation in 100% EtOH for 3*1h (3*3 min for thin sections), 96% EtOH for 2*1h (2*2 min for thin sections), and 70% EtOH for 1h (2 min for thin sections) and then washed in PBS for 10 min (2*1 min for thin sections).

### Optical Clearing

Samples were prepared according to a slightly modified version of a previously published fast and simple protocol^9^. Samples were sectioned to 200 μm sections using a Vibratome and then de-lipidated in clearing solution for 2 h at 70-80°C (depending on age and species, has to be optimized for each application). For thin sections, sections were gently removed from glass slides using a razor blade, and then cleared for 1h at 80°C free-floating in an Eppendorf tube.

### Labelling

PBST was used as antibody diluent for mouse samples and HEPES-TCS buffer (10 mM HEPES pH7.5 with 200 mM NaCL and 10% TritonX-100) was used for human samples. Samples were incubated in primary antibody at 37°C for 2-24 hours, washed once in antibody diluent for minimum 5 minutes, then incubated in secondary antibody at 37°C for 2-24 hours. Samples were then washed in antibody diluent for minimum 5 minutes before mounting. Antibodies used were a rabbit polyclonal antibody to podocin (Sigma-Aldrich, P0372) used at a dilution of 1:100, sheep polyclonal antibody to nephrin (R&D systems, AF4269) used at a dilution of 1:50 and a rabbit polyclonal antibody to collagen IV (Abcam, ab256353). Secondary antibodies used were a donkey anti-rabbit (Jackson Immunosearch, 711-005-152) conjugated to Atto-594 and a donkey anti-sheep (Jackson Immunosearch, 713-005-147) conjugated to either Alexa-488 (human samples) or Abberior STAR635P (mouse samples). The secondary antibodies were conjugated to dyes in house as previously described^9^. The collagen IV primary antibody above was directly conjugated to Alexa-555 in-house as previously described^9^.

For thin sections, PBST supplemented with 3% Bovine Serum Albumin and 5% normal donkey serum was used as antibody diluent and PBST was used for washing steps. After de-paraffinization, a sheep anti-nephrin and a rabbit anti-podocin (same antibodies as above) antibody was added at a 1:200 and 1:400 dilution, respectively, and incubated at room temperature for 3 hours. After this, sections were washed 5*2 min at room temperature. A donkey anti-rabbit Atto-594 and a donkey anti-sheep Abberior STAR635P antibody was added, both at a 1:400 dilution and incubated at 3 hours at room temperature.

### Mounting

Samples were incubated in 80% wt/wt fructose (1 mL of dH20 added to 4g of fructose, 0.5% 1-Thioglycerol was added to inhibit the Maillard reaction) with (human samples) or without (mouse samples) 4M Urea at 37°C with shaking at 500 rpm for 15 minutes and then placed in a MatTek or Ibidi dish along with a few drops of fructose solution with a cover slip on top (to prevent evaporation) prior to imaging. Thin sections on slides were mounted in a drop of 80% wt/wt fructose solution (see above).

### Imaging

Images were acquired using a Leica SP8 STED 3X system. Objectives and other relevant imaging parameters (such as pinhole diameter) is indicated in figure legends.

### Image Processing and Analysis

Images were smoothened (unless other stated) by replacing each pixel value by the mean of its 3×3 neighborhood using Fiji/ImageJ before inclusion in figures. Details regarding the ImageJ macro and extracted morphometric parameters can be found elsewhere^7,8^.

### Schemes

The scheme in Fig. 1a was produced by D.U-J using BioRender.

## Acknowledgements

This study was supported by the Clinical Research Unit (CRU) 329 of the Deutsche Forschungsgemeinschaft (DFG) and partly by FOR 2743 of DFG as well as by Else Kröner-Fresenius-Stiftung, by the German Research Foundation under Germany’s Excellence Strategy - EXC 2030: CECAD - Excellent in Aging Research - Project number 390661388 and by the Koeln Fortune program / Faculty of Medicine, University of Cologne outside the conduct of this project. We thank the CECAD Imaging Facility for their support in the acquisition of microscopy data.

