## Supplementary Figures for "Three-dimensional Super-resolved Imaging of Paraffin-embedded Kidney Samples"

**Running title:** Fast Protocol for FFPE Kidney Imaging

**Corresponding author:**
David Unnersjö-Jess
Uniklinik Köln
Kerpener Strasse 62
509 37 Köln
Germany

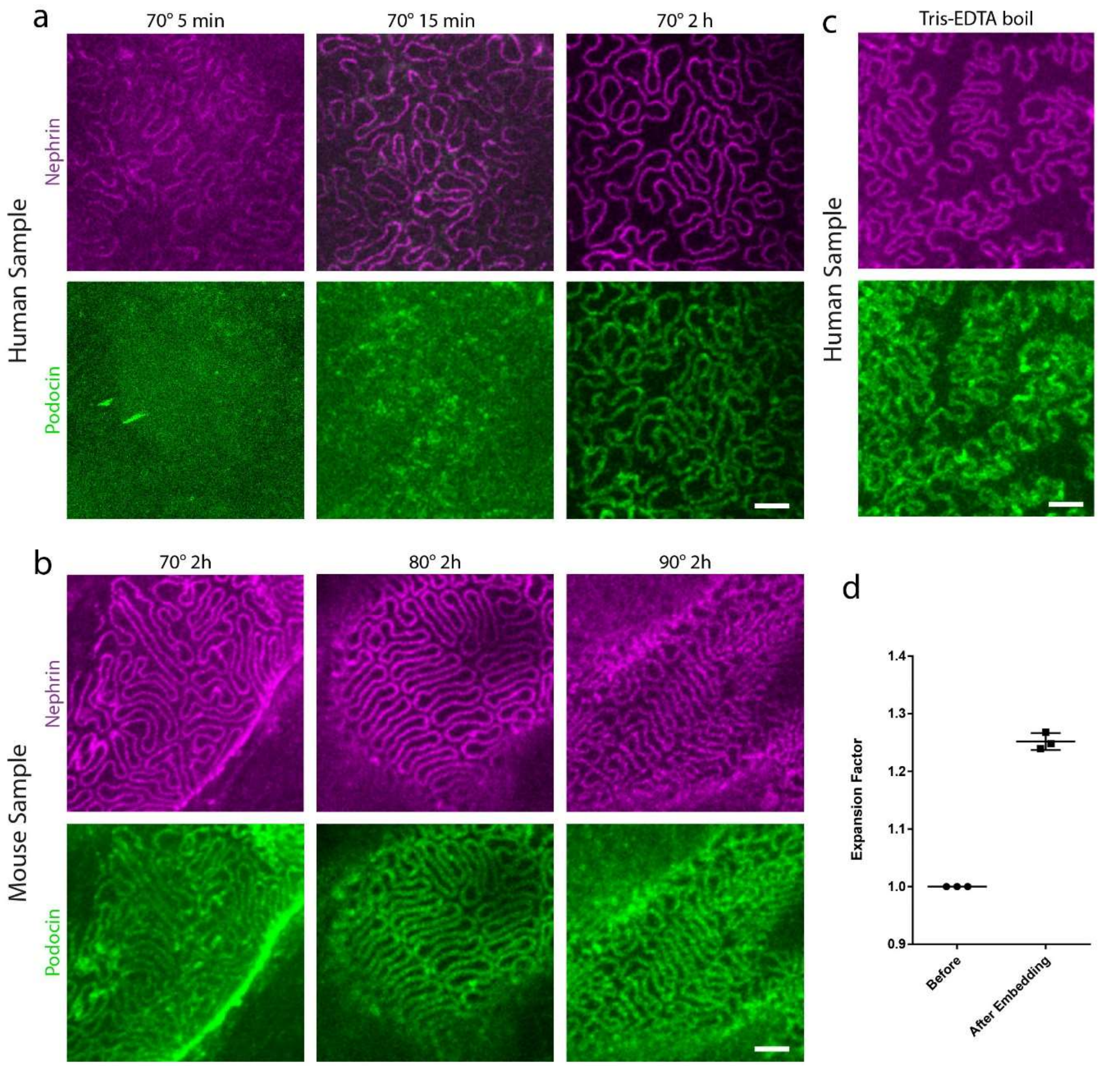


**Supplementary Figure S1.** Effect of different times and temperatures of the SDS de-lipidation step as evaluated by STED imaging of the two slit diaphragm proteins nephrin (labelled with Abberior STAR-635P) and podocin (labelled with Atto-594). Scale bars 1 µm. (a) SDS de-lipidation of a human control sample (nephrectomy, renal carcinoma) at 70°C for different duration as indicated. Note that the nephrin staining was of sufficient quality after just 15 min of clearing whereas the podocin staining required 2 hours of de-lipidation for optimal results. (b) SDS de-lipidation of a mouse wild-type sample for 2 hours at different temperatures as indicated. The 2 hours at 70°C which was sufficient for the human sample was not sufficient for the mouse sample, but increasing the temperature to 80°C resulted in sufficient staining quality. At 90°C, nanoscale distortions could be observed. (c) The same human control sample as in (a) treated with a standard Tris-EDTA boil at 95°C for 30 min showed higher background and the podocin staining appeared slightly smeared out and distorted. (d) Expansion factor of the protocol. The perimeter of three different mouse kidney tissue samples were measured using ImageJ. Measurements were performed prior to SDS clearing (after de-paraffinization) and after embedding in 80% fructose with 4M Urea. The expansion factor was taken as the perimeter after embedding divided by the perimeter prior to SDS clearing. Each dot represents one sample, the black line represents mean and the error bars represent standard deviation.

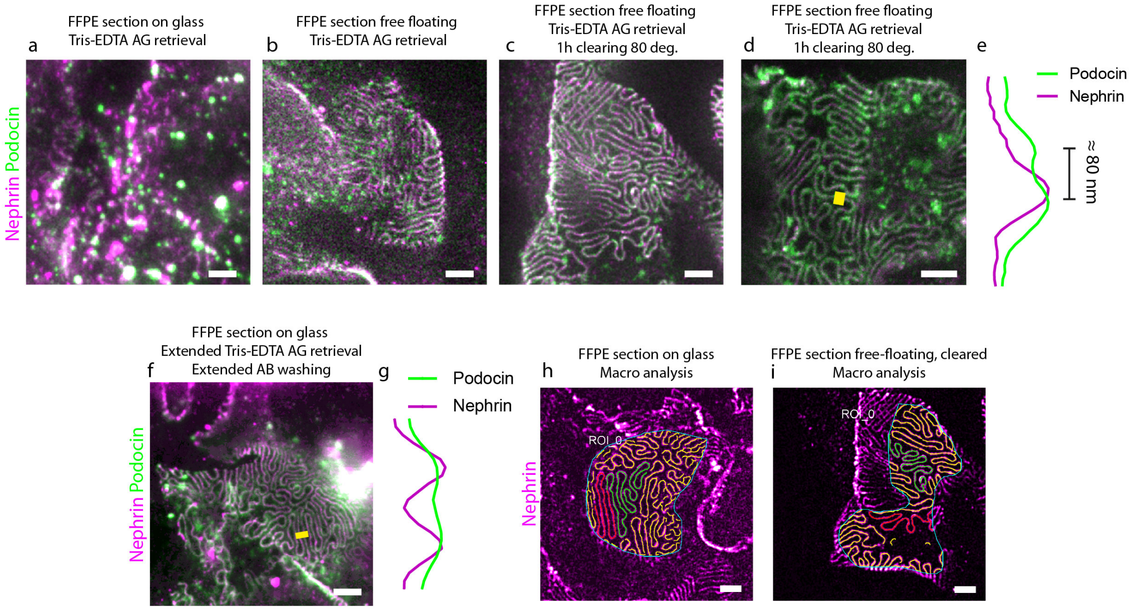


**Supplementary Figure S2.** Performance of our protocol on 10 um paraffin sections. Samples were all stained for nephrin with Abberior STAR635P and for podocin with Atto594 and images using a 100X NA 1.4 oil objective. Scale bars 1 µm. (a) Representative STED image when using a standard Tris-EDTA antigen retrieval for 20 min. We applied the same simplified staining protocol as we used for cleared sections. A high degree of unspecific staining is observed and FP can barely be visualized. (b) By applying the same AG retrieval and staining protocol as in (a) but then removing the section from the glass and performing the antibody incubation in a free-floating manner increased staining quality substantially, and foot processes can be resolved. (c) It is only when applying an SDS de-lipidation step to free-floating sections that the highest staining quality and contrast is achieved. (d) In some places, podocin could be resolved on both sides of the slit diaphragm only in the de-lipidated sample. (e) Line plot of the yellow line indicated in (d) showing the two podocin lines with a separation of around 80 nm. (f) By increasing the Tris-EDTA AG retrieval time to 40 minutes, and by applying an extended staining protocol with BSA/NDS-blocking and several washing steps between incubations, STED imaging could be performed also on sections still mounted on glass. (g) Line plot of the yellow line indicated in (f). The two podocin lines could not be resolved in these samples, probably as a result of the inhibited spatial expansion of the sample when mounted on a glass. (h-i) Results of the SD segmentation ImageJ macro on STED images of FFPE sections mounted on glass (h) and free-floating and cleared (i).

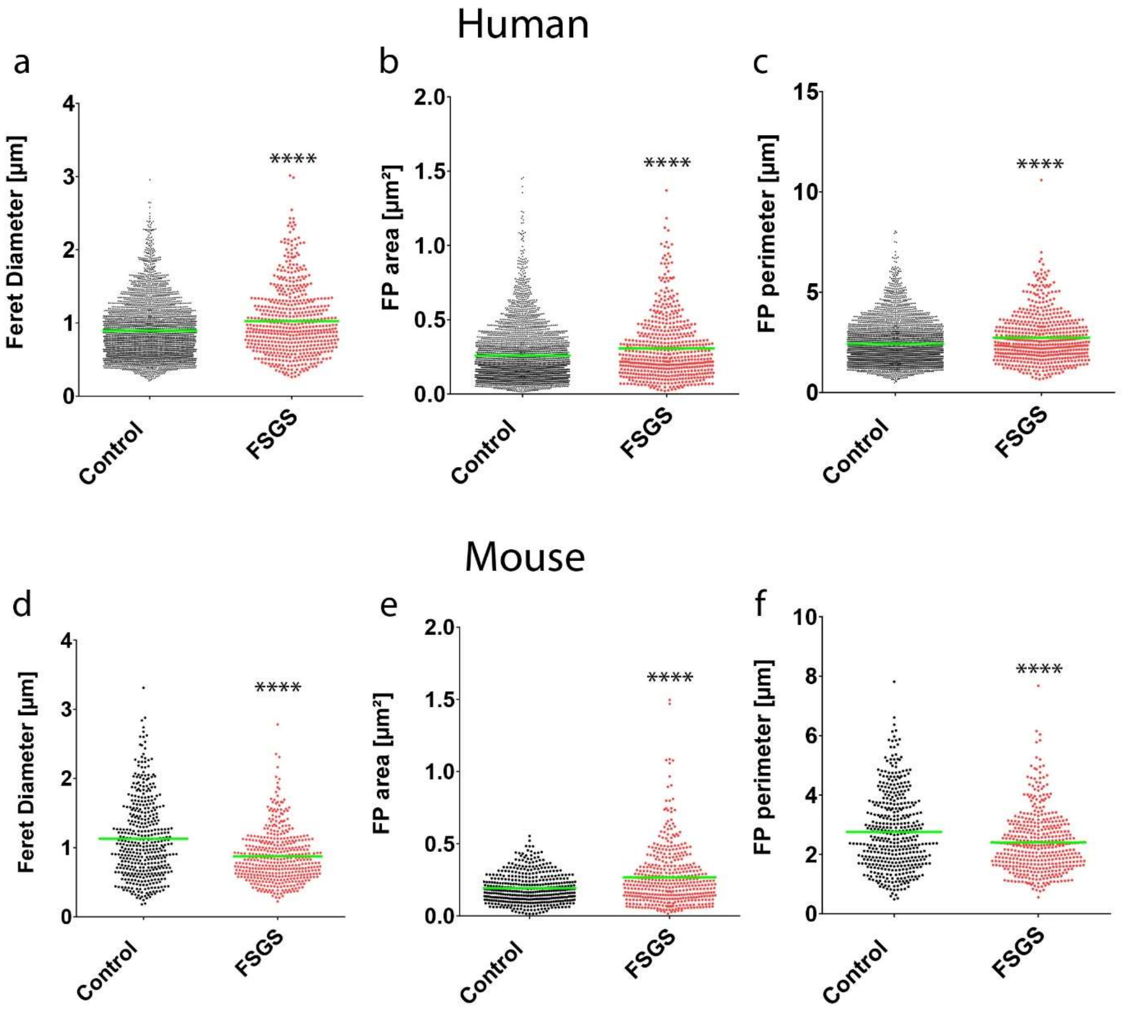


**Supplementary Figure S3.** Further morphometric parameters extracted from the patients/mice in Fig. 3. (a-c) Feret diameter (a), FP area (b) and FP perimeter (c) for control and FSGS patients. Each dot represents one FP. Green line represents mean. P-values < 0.0001 for all parameters. (d-f) Feret diameter (d), FP area (e) and FP perimeter (f) for control and FSGS mice. Each dot represents one FP. Green line represents mean. P-values < 0.0001 for all parameters.

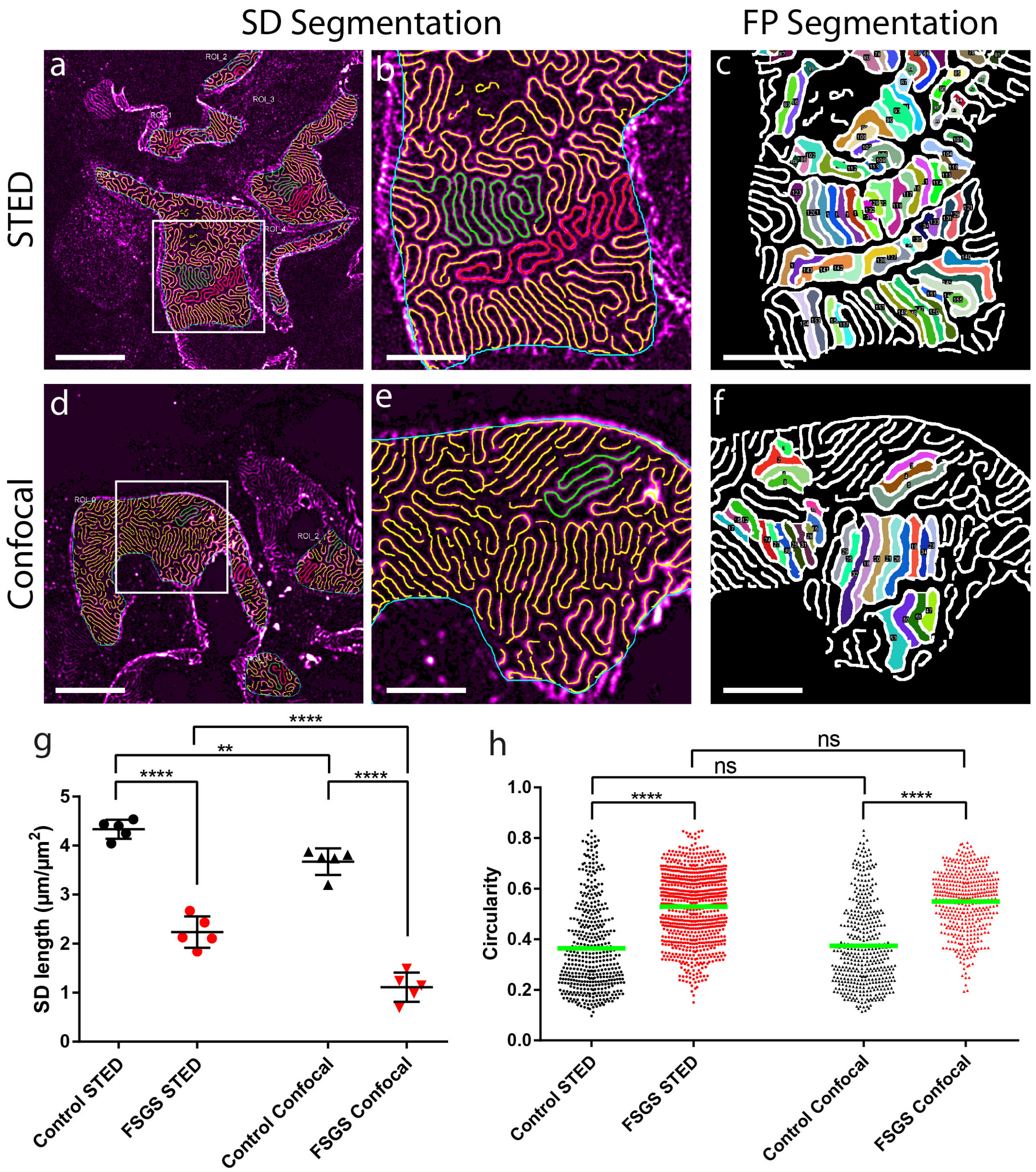


**Supplementary Figure S4.** Representative comparison of performance of the ImageJ macro-based analysis on images acquired with confocal and STED microscopy. (a) STED image of foot processes in a WT mouse with result of the SD segmentation macro in overlay. Scale bars 5 µm (a,d) and 2 µm (b-c, e-f). (b) Zoom of the area indicated by a white square in (a). (c) Result of the FP segmentation macro in the same image as in (b). (d) Confocal image of foot processes in a WT mouse with result of the SD segmentation macro in overlay. (e) Zoom of the area indicated by a white square in (d). Occasional failure of the macro to segment the SD can be observed. (f) Result of the FP segmentation macro in the same image as in (e). Since this result is based on the SD segmentation in (e), the number of foot processes that are segmented from an image is lower. (g-h) Quantification of SD length per area (g) and circularity score (h) for control and mutant (FSGS) mice in STED versus confocal images as indicated in plots. In (g), each dot represents one image, black line represents mean and the error bars represent standard deviation. In (h), each dot represents one FP and the green line represents mean. One-way ANOVA was used to determine statistical significance. ** $p < 0.01$, **** $p < 0.0001$. In both WT and FSGS mice, a lower SD length is extracted from confocal images as compared to STED images, which is a consequence of the slightly lower segmentation accuracy in confocal images (b, e). No significant difference is seen for the FP circularity between STED and confocal since this parameter is only dependent on the shape of each segmented foot process. The lower number of segmented FP is reflected in the lower number of data points for confocal images in (h).

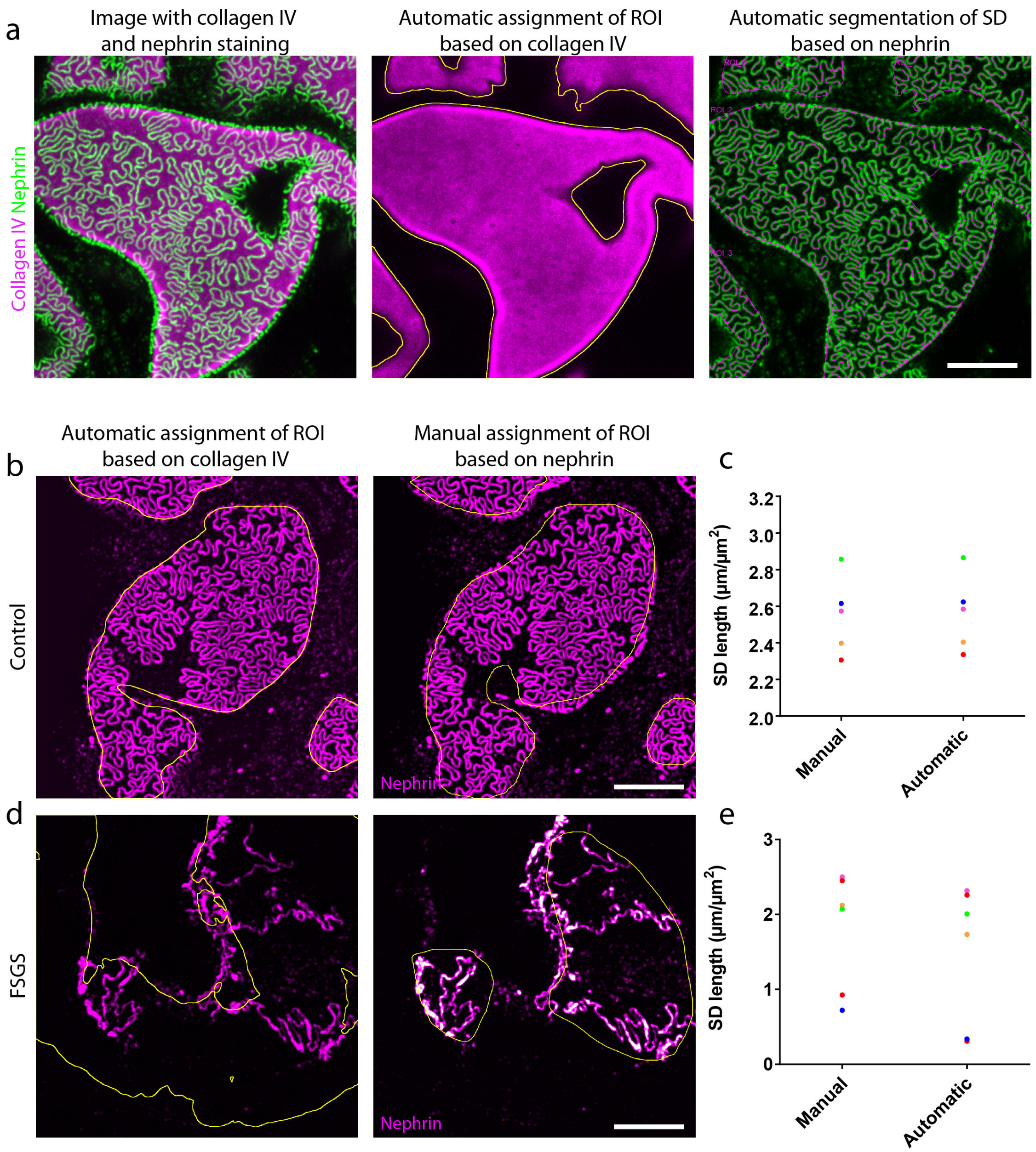


**Supplementary Figure S5.** Automatic assignment of ROI for analyzing SD length with the ImageJ macro based on a collagenIV staining. All samples were stained for Nephrin using Alexa-488 and for collagenIV using Alexa-555 and imaged using a 100X NA 1.4 oil objective with a pinhole setting of 0.3 AU. Samples are the same human samples as in Fig. 2-3. Scale bars 5 µm. (a) Representative result of the automatic segmentation of capillary surface by staining for the glomerular basement membrane using a collagenIV antibody. (b-d) Comparison between automatic (collagenIV-based) and manual assignment of the ROI in samples from a control (b) and FSGS (d) patient. (c) and (e) shows the resulting SD length extracted with the macro based on the two ROI assignment strategies. Each dot represents one image and each color represents one specific image. The manual assignment is feasible in a control sample where the SD pattern is dense, but on capillaries with severe effacement (d) the manual assignment leads to a slight over-estimation of the SD density, due to an under-estimation of the capillary surface.

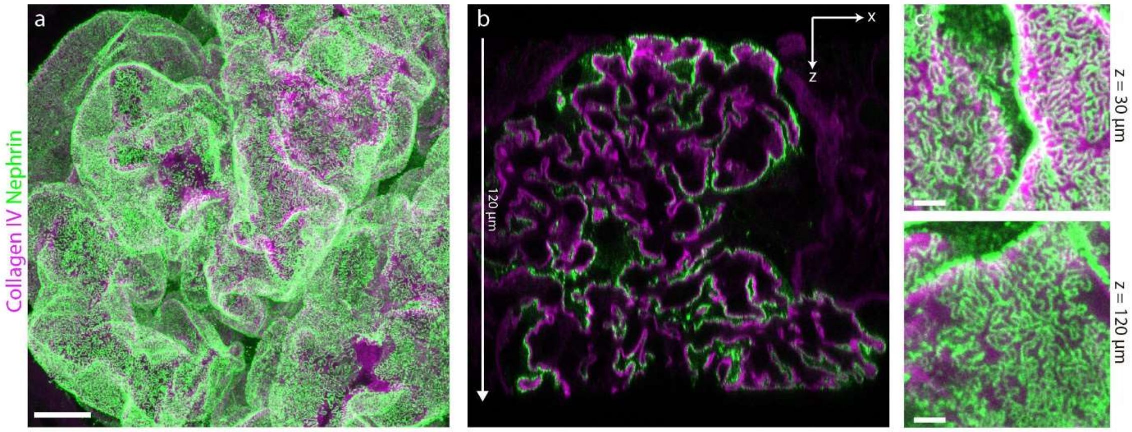


**Supplementary Figure S6.** Three-dimensional confocal imaging of a whole human glomerulus at imaging depths > 100 µm. Sample was stained for collagen-IV with Alexa-555 and nephrin using Alexa-488 and imaged using a 93X NA 1.3 glycerol motcorr objective. (a) x-y view of a maxiumum intensity projection of a 120 µm thick z-stack from the human control sample. (b) x-z slice of the same stack as in (a). (c) Maximum intensity-projected sub-stacks of around 10 µm in thickness acquired at the indicated depth shows that foot processes can be resolved throughout the dataset in (a-b). Scale bars 10 µm (a), 2 µm (c).
